# Strengthening of the efferent olivocochlear system leads to synaptic dysfunction and tonotopy disruption of a central auditory nucleus

**DOI:** 10.1101/433581

**Authors:** Mariano N. Di Guilmi, Luis E. Boero, Valeria C. Castagna, Adrián Rodríguez-Contreras, Carolina Wedemeyer, María Eugenia Gómez-Casati, Ana Belén Elgoyhen

**Affiliations:** Instituto de Investigaciones en Ingeniería Genética y Biología Molecular, Dr. Héctor N Torres, INGEBI-CONICET, Buenos Aires, Argentina. (C1428ADN); Instituto de Farmacología, Facultad de Medicina, UBA, Buenos Aires, Argentina. (C1121ABG); Department of Biology, the City College of the City University of New York, New York, NY, USA. (NY 10031)

**Keywords:** α9α10 nAChR, Chrna9L9’T, efferent MOC inhibition, tonotopy, MNTB

## Abstract

The auditory system in many mammals is immature at birth but precisely organized in adults. Spontaneous activity in the inner ear plays a critical role in guiding this process. This is shaped by an efferent pathway that descends from the brainstem and makes transient direct synaptic contacts with inner hair cells (IHCs). In this work, we used an α9 cholinergic receptor knock-in mouse model (of either sex) with enhanced medial efferent activity (*Chrna9L9’T*, *L9’T*) to understand the role of the olivocochlear system in the correct establishment of auditory circuits. Wave III of auditory brainstem responses (which represents synchronized activity of synapses within the superior olivary complex) were smaller in *L9’T* mice, suggesting a central dysfunction. The mechanism underlying this functional alteration was analysed in brain slices containing the medial nucleus of the trapezoid body (MNTB), where neurons are topographically organized along a medio-lateral axis. The topographic organization of MNTB physiological properties observed in WT mice was abolished in the *L9’T* mice. Additionally, electrophysiological recordings in slices evidenced MNTB synaptic alterations, which were further supported by morphological alterations. The present results suggest that the transient cochlear efferent innervation to IHCs during the critical period before the onset of hearing is involved in the refinement of topographic maps as well as in setting the correct synaptic transmission at central auditory nuclei.

**Significance Statement:** Cochlear inner hair cells of altricial mammals display spontaneous electrical activity before hearing onset. The pattern and firing rate of these cells is crucial for the correct maturation of the central auditory pathway. A descending efferent innervation from the central nervous system contacts hair cells during this developmental window. The function of this transient efferent innervation remains an open question. The present work shows that the genetic enhancement of efferent function disrupts the orderly topographic distribution at the medial nucleus of the trapezoid body level and causes severe synaptic dysfunction. Thus, the transient efferent innervation to the cochlea is necessary for the correct establishment of the central auditory circuitry.

## Introduction

The strength and physiological properties of synaptic inputs, the accurate organization of neuronal circuits and the formation of topographic maps in the mature brain are established during development, through activity-dependent processes that involve reorganization and fine tuning of immature synaptic and cellular networks (Goodman and Shatz, 1993; Hanson and Landmesser, 2004; Kirkby et al., 2013). The auditory system in many mammals is very immature at birth but precisely organized in adults. Spontaneous activity in the inner ear during an early developmental critical period comes into play to guide this process (Lippe, 1994; Kotak and Sanes, 1995; Jones et al., 2007; Tritsch et al., 2007; Sonntag et al., 2009). Spontaneous activity, driven by calcium action potentials in IHCs (Kros et al., 1998; Glowatzki and Fuchs, 2000; Marcotti et al., 2003; Tritsch et al., 2007; Johnson et al., 2011), is essential for several processes related to the survival of target neurons in the cochlear nucleus (Leake et al., 2006), accurate wiring of auditory pathways (Friauf and Lohmann, 1999) and the refinement of tonotopic maps in the lateral superior olive (Kandler, 2004; Clause et al., 2014).

A distinctive feature of IHCs during the prehearing critical period is the presence of direct axo-somatic efferent medial olivocochlear (MOC) synaptic contacts, which disappear at hearing onset when finding their final targets, the outer hair cells (Warr and Guinan, 1979; Simmons et al., 1996). This synapse is cholinergic (Glowatzki and Fuchs 2000; Katz et al., 2004, Gomez-Casati et al., 2005) and mediated by highly calcium permeable α9α10 nicotinic cholinergic receptors (nAChRs) present in IHCs (Elgoyhen 1994, 2001; Weisstaub et al., 2002; Lipovsek et al., 2012), coupled to the activation of small conductance, calcium activated SK2 potassium channels (Glowatzki and Fuchs, 2000). Exogenously applied acetylcholine (Glowatzki and Fuchs, 2000) or electrical stimulation of efferent terminals (Goutman et al., 2005; Wedemeyer et al., 2018) inhibits IHC action potentials. Therefore, it has been proposed that cholinergic efferent inhibition of IHCs might impose rhythmicity onto IHC action potential generation and spontaneous activity of the auditory pathway during the critical developmental period preceding hearing onset (Glowatzki and Fuchs, 2000; Johnson et al., 2011; Sendin et al, 2014; Moglie et al., 2018). However, this notion has been challenged (Tritsch et al., 2010a), and therefore the function of the developmental efferent innervation is still a matter of debate.

In a recent paper Clause et al., (2014) showed that in α9 knock-out mice, which lack efferent activity (Vetter et al., 1999; 2007), the spike patterning of spontaneous activity at the MNTB level is altered, leading to a reduced permanent sharpening of functional topography (Clause et al., 2014) and impairment of sound localization and frequency processing (Clause et al., 2017). The present work shows an alternative approach, in which we used an α9 knock-in mouse model (*L9’T*) with enhanced efferent activity (Taranda et al., 2009) leading to sustained inhibition of IHC action potential generation (Wedemeyer et al., 2018). Since a decrease in efferent activity leads to alteration in functional topography, one could *a priori* hypothesize that the enhancement of activity might lead to hyper refinement of topographic properties. To this end we analyzed the calyx of Held - MNTB synapse, making focus on the MNTB innervation, the synaptic features and its topography. Synaptic transmission and synaptic structure of the calyx of Held was greatly impaired in α9 *L9’T* mice. The presence of multiple innervation to the calyx was evidenced at a developmental age where this feature should be absent or considerably reduced. An electrophysiological correlate was observed since the proportion of immature “small”-evoked postsynaptic currents (EPSCs) was significantly enhanced in *L9’T* mice. Contrary to our hypothesis, a complete lack of topographic organization of the MNTB was observed. Taken together, these results provide clear evidence that a tight regulation of pre-hearing spontaneous activity is key for the development of the central auditory pathway, and that this is brought about by the transient MOC innervation to IHCs.

## Methods

### Animals and experiments

Generation of the knock-in mouse (*L9’T*) has been described previously (Taranda et al., 2009). Wild-type (WT) or homozygous *L9’T* mice of either sex were used. All experimental protocols were carried out in accordance with the American Veterinary Medical Associations’ AVMA Guidelines on Euthanasia (2013) and approved by the IACUC at INGEBI.

### Auditory brainstem responses (ABRs)

Animals of postnatal days (P) 14, 16 and 21 were anesthetized with a mix of xylazine (10 mg/kg i.p.) / ketamine (100 mg/kg i.p) and needle electrodes were inserted at vertex and pinna, with a ground near the tail. ABRs were evoked with 5 ms tone pips (0.5 ms rise-fall, cos2 onset, at 35/s). The response was amplified (10000X) filtered (0.1-3 kHz) and averaged with an A-D board in a LabVIEW-driven data-acquisition system (National Instruments, Austin, Texas, USA, RRID:SCR_014325). Sound level was raised in 5 dB steps from 20 dB to 80 dB SPL. At each level, 1024 responses were averaged (with stimulus polarity alternated), using an “artefact reject” whereby response waveforms were discarded when peak-to-peak amplitude exceeded 15 μV. Upon visual inspection of stacked waveforms, “threshold” was defined as the lowest SPL level at which a wave could be detected. Wave I, II and III were identified and the peak to peak amplitude computed by off-line analysis of stored waveforms using Clampfit 10.3 (Molecular Devices, LLC, San José, CA, USA).

### Electrophysiology on MNTB slices

For slice recordings, 50 mice of either sex between 12 and 14 postnatal days old were used. Their brains were removed rapidly after decapitation and placed into an ice-cold low-Ca^2+^ artificial cerebrospinal fluid solution (aCSF). This solution contained the following (in mM): 125 NaCl, 2.5 KCl, 3 MgCl_2_, 0.1 CaCl_2_, 1.25 NaH_2_PO_4_, 0.4 ascorbic acid, 3 myoinositol, 2 pyruvic acid, 25 D-glucose, and 25 NaHCO_3_. The brainstem was glued on a cooled chamber of a vibrating microslicer (Vibratome 1000 Plus, Ted Pella, California, USA). Transverse slices (300 μm thickness) containing the MNTB were sequentially cut and transferred into an incubation chamber containing normal aCSF at 37°C for 30 min. After incubation, slices were allowed to return to room temperature. Normal aCSF had the same composition as the slicing solution except that the MgCl_2_ and CaCl_2_ concentrations were 1 and 2 mM, respectively. The pH was 7.4 when gassed with 95% O_2_ and 5% CO_2_.

#### Whole-cell patch-clamp recordings

Slices were transferred to an experimental chamber. During recording, slices were continuously perfused with carbogenated (95% O_2_ and 5% CO_2_) aCSF maintained at room temperature (22–25°C). Medial (M) and lateral (L) MNTB principal cells were compared in order to evaluate topographic differences. The position of these two opposite groups of cells was determined during the experiment according to the morphological landmark delimited by the middle line (M) and the VII cranial nerve (L). Lucifer Yellow (0.3 mg/ml; Sigma Aldrich) was added to the internal solution to certify the position at the end of the experiment. MNTB neurons were visualized using a Zeiss Axioskop microscope (Oberkochen, Germany) and viewed with differential interference contrast by a 40X water-immersion objective (0.8 numerical aperture water-immersion objective) and a camera with contrast enhancement (DMK 23UP1300, The Imaging Source, North Carolina, USA). Whole-cell recordings were made with patch pipettes pulled from thin-walled borosilicate glass (World Precision Instruments, Florida, USA, RRID:SCR_008593). Electrodes had resistances of 3.8 to 4.5 MΩ. Potassium currents were isolated by an internal solution containing (mM): k-gluconate 121.3, KCl 20, Hepes 10, phosphocreatine 10, EGTA 0.5, Mg-ATP 4, Li-GTP 0.3. Additionally, external solution was supplemented with TTX (1 μm), CdCl_2_ (50 μm) and ZD7288 (10 μm). Hyperpolarization-activated currents (I_h_-type) were analysed from recordings in presence of TTX (1 μM) and TEA (20 mM). The pH was adjusted to 7.3 with KOH. The pipette solution used for isolating synaptic currents was (in mM): CsMeSO_3_ 135, TEA Cl 13, HEPES 5, MgCl_2_ 3.5, CaCl_2_ 0.1, Na_2_-ATP 2.5, EGTA 1; pH 7.2 (CsOH). Liquid junction potential was uncompensated in both solutions.

Patch clamp recordings were made using an Axopatch 200A (Molecular Devices, San Jose, CA, USA) amplifier, a Digidata 1320 (Molecular Devices, San Jose, CA, USA) and pClamp 9.0 software (Molecular Devices, San Jose, CA, USA; RRID:SCR_011323). Data were sampled at 50 kHz and filtered at 4-6 kHz (low pass Bessel). Series resistances ranged from 6 to 15 MΩ. Whole-cell membrane capacitance (15-25 pF) was registered from the amplifier after compensation of the transient generated by a 10 ms voltage step. To elicit action potentials, positive current steps (100-500 pA) during 500 ms were applied. Voltage-clamp protocols were as follows: the cell was held at a resting potential of −50 mV (1 sec) and stepped to a range of −130 mV to −40 mV for 500 ms (in increments of 5 mV). I_h_ amplitude was measured as instantaneous (I_I_) minus slowly inward (I_s_) current (I_S_ − I_I_; Yi et al., 2010). Statistical comparisons were evaluated at its maximal current amplitude (−140 mV). ZD7288 (50 μM) or CsCl (1 mM) were used to confirm the presence of a I_h_ current. No leak current was subtracted from any of the raw current traces for I_h_ current. Potassium currents were elicited by square depolarizing pulses of 1s from −90 to +40 mV. Off-line leak subtraction was performed for K^+^ currents and the average current at the steady-state (within the last 50 ms) is reported.

Miniature excitatory post synaptic currents (mEPSCs) were recorded continuously for at least three separate periods of 1 min. Amplitude and frequency were analysed using Clampfit 10.3 (Molecular Devices, San Jose, CA, USA) and Mini Analysis Program (Synaptosoft, Decatur, GA, RRID:SCR_002184). EPSCs were evoked by stimulating the globular bushy cell axons in the trapezoid body at the midline using a bipolar platinum electrode and an isolated stimulator (0.1 ms duration and 50-200 μA amplitude). Strychnine (1 μM) was added to the aCSF to block inhibitory glycinergic synaptic responses. Data distribution was evaluated by fitting with a gaussian function (extreme value distribution) with the Statistica Software (Stat Soft. Inc, Tulsa, Ok, RRID:SCR_014213). This distribution allows to evidence the largest extreme.

#### Analysis of electrophysiological data

Electrophysiological recordings were analysed using Clampfit 10.3 (Molecular Devices, San Jose, CA, USA) and custom routines in Igor 6.2 (Wavemetrics, Portland, OR, USA, RRID:SCR_000325). Igor routines were used to perform offline leak subtraction (for potassium currents). Series resistances of EPSC recordings was ~ 50-70% compensated. The remaining Rs error of postsynaptic currents was corrected by an off-line Igor routine.

### Morphological analysis

*In-vivo* dye injections were performed to identify individual axons and calyx-like terminals contacting a principal MNTB neuron. For *in-vivo* tracing, a ventral craniotomy was performed (Rodriguez-Contreras et al., 2008) and afferent fibres to the calyx of Held were electroporated with rhodamine-labeled dextran (3,000 MW; D3308 - Invitrogen, Carlsbad, California, USA) or microruby (3,000 MW, D7162 - Invitrogen, Carlsbad, CA, USA) at the midline, in pups between P12 and P14. Thirty minutes later, the animal was perfused with parafomaldehyde (4%), and the brainstem was sliced with a cryostat into 60 μm-thick sections. Brainstem sections were mounted with a medium containing DAPI for nucleus identification (Vectashield Antifade Mounting Medium with DAPI; Vectorlabs, Burlingame, CA, USA). A laser scanning confocal microscope (Zeiss LSM 800 or Leica TCS SPE) equipped with krypton-argon and helium-neon lasers was used to acquire high resolution z-stack images (40X, NA 1.3 oil, 1.14 μm steps; 63X, NA 1.4 oil, 0.5 μm steps) of randomly lateral or medial sections of the MNTB. Images were opened and analysed with Fiji software (Schindelin et al., 2012). Digitized tracing for each axonal profile was made with Fiji.

### Experimental Design and Statistical Analysis

Experiments were designed in order to reduce the number of animals but taking into account a balance between the number of samples to accurately perform statistical tests and the ethics guidelines for animal research as described above. All statistical tests were carried out with Statistica 7.0 software (Stat Soft. Inc, Tulsa, OK, USA, RRID:SCR_014213) with the exception of ABRs performed with Prism 6 software (GraphPad, La Jolla, CA, USA, RRID: 294 SCR_002798). Prior to performing any analysis, data sets were tested for normal distribution and homoscedasticity. If these assumptions were satisfactorily passed, a parametric test was applied. In these cases, comparisons were made by one-way ANOVA and the statistic “F” value with the associated “p-value” significance was reported in every case. Otherwise, non-parametric Mann-Whitney test was used. Values of p<0.05 were considered significant. Average data were expressed and plot as mean ± S.E.M. In all cases “n” indicates the number of cells tested.

### Drugs and reagents

All drugs and reagents were purchased from Sigma-Aldrich (Saint Louis, Missouri, USA, RRID:SCR_008988) with the exception of ZD7288 and TTX which were purchased from Tocris Bioscience (Bristol, UK; RRID: SCR_003689).

## Results

### Reduction of ABR wave III amplitude in *L9’T* mice

In *L9’T* mice the MOC-IHC synapse displays both pre- and postsynaptic alterations leading to an enhancement and prolongation of inhibitory synaptic responses (Wedemeyer et al., 2018). Furthermore, low-frequency stimulation of MOC fibers allows a complete suppression of IHC action potentials (Wedemeyer et al., 2018). In order to examine whether this enhanced MOC efferent cholinergic activity has an impact on the functionality of the auditory pathway, auditory brainstem responses (ABRs) were measured at P16 (temporally close to the onset of hearing, Table 1) and at P21 (considered as an auditory mature stage, Fig. 1A and Table 1). ABR individual waves reflect the activation of subsequent auditory processing stations and are used as a functional hearing test that detects retro-cochlear abnormalities underlying hearing impairment (Karplus et al., 1988; Shapiro, 1988; Shaw, 1988). Pip-evoked ABR waveform amplitudes were analyzed at 80 dB SPL for peaks corresponding to the auditory nerve (wave I), cochlear nucleus (wave II) and superior olivary complex (SOC) (wave III) at different frequencies (8, 16 and 32 KHz) (Melcher et al., 1996) (Fig. 1A). As recently shown (Boero et al., 2018), no abnormalities in peak I amplitude were observed in *L9’T* compared to WT mice at 8, 16 and 32 KHz either at P16 (Table 1) or P21 (Fig. 1B; Table 1). This suggests no gross alterations in the first synapse of the auditory pathway between IHCs and auditory nerve fibers in *L9’T* mice. The amplitude of peak II which is the second neural relay, the cochlear nucleus, was diminished at P16 in mice with enhanced MOC activity compared to WT at 8 KHz, 16 KHz and 32 KHz (Table 1). However, this difference disappeared in the auditory mature stage (Table 1 and Fig. 1C). Interestingly, peak III amplitude at P16 was reduced in *L9’T* mice compared to WT at 8KHz and at all frequencies tested at P21 (Table 1 and Fig 1D). These results show that a cochlear enhancement of α9α10 nAChR activity results in a central dysfunction at the brainstem level.

**Table 1.** ABR quantification at P16 and P21. ^1^p = 0.05; ^2^p = 0.027; ^3^p = 0.023; ^4^p = 0.035; ^5^p = 0.031; ^6^p = 0.0032; ^7^p = 0.0026. One-way ANOVA

|  |  | Peak I (μV) |  | Peak II (μV) |  | Peak III (μV) |  |
| --- | --- | --- | --- | --- | --- | --- | --- |
|  |  | WT | *L9'T* | WT | *L9'T* | WT | *L9'T* |
| P16 | 8 kHz | 0.57±0.05 (n=10) | 0.39±0.08 (n=13) | 0.28±0.04 (n=10) | 0.17±0.04 (n=13)<sup>1</sup> | 0.28±0.05 (n=10) | 0.16±0.03 (n=13)<sup>4</sup> |
|  | 16 kHz | 0.47±0.05 (n=9) | 0.44±0.07 (n=12) | 0.48±0.06 (n=9) | 0.31±0.04 (n=12)<sup>2</sup> | 0.39±0.08 (n=9) | 0.36±0.04 (n=12) |
|  | 32 kHz | 0.44±0.04 (n=9) | 0.44±0.06 (n=12) | 0.47±0.07 (n=9) | 0.26±0.05 (n=12)<sup>3</sup> | 0.31±0.07 (n=9) | 0.25±0.04 (n=12) |
| P21 | 8 kHz | 0.62±0.05 (n=24) | 0.61±0.06 (n=22) | 1.39±0.13 (n=24) | 1.37±0.09 (n=22) | 1.19±0.10 (n=24) | 0.90±0.08 (n=22)<sup>5</sup> |
|  | 16 kHz | 0.51±0.04 (n=25) | 0.55±0.04 (n=22) | 1.22±0.07 (n=24) | 1.21±0.08 (n=22) | 1.13±0.07 (n=25) | 0.86±0.05 (n=22)<sup>6</sup> |
|  | 32 kHz | 0.52±0.04 (n=24) | 0.50±0.03 (n=22) | 0.86±0.08 (n=24) | 0.76±0.06 (n=22) | 0.74±0.06 (n=24) | 0.74±0.06 (n=22)<sup>7</sup> |

<sup>1</sup>F:3.55, p=0.05; <sup>2</sup>F:5.78, p=0.027; <sup>3</sup>F:6.07, p=0.023; <sup>4</sup>F:5.10, p=0.035; <sup>5</sup>F:5.02, p=0.031; <sup>6</sup>F:9.7, p=0.0032; <sup>7</sup>F:10.22, p=0.0026

**Figure 1.**
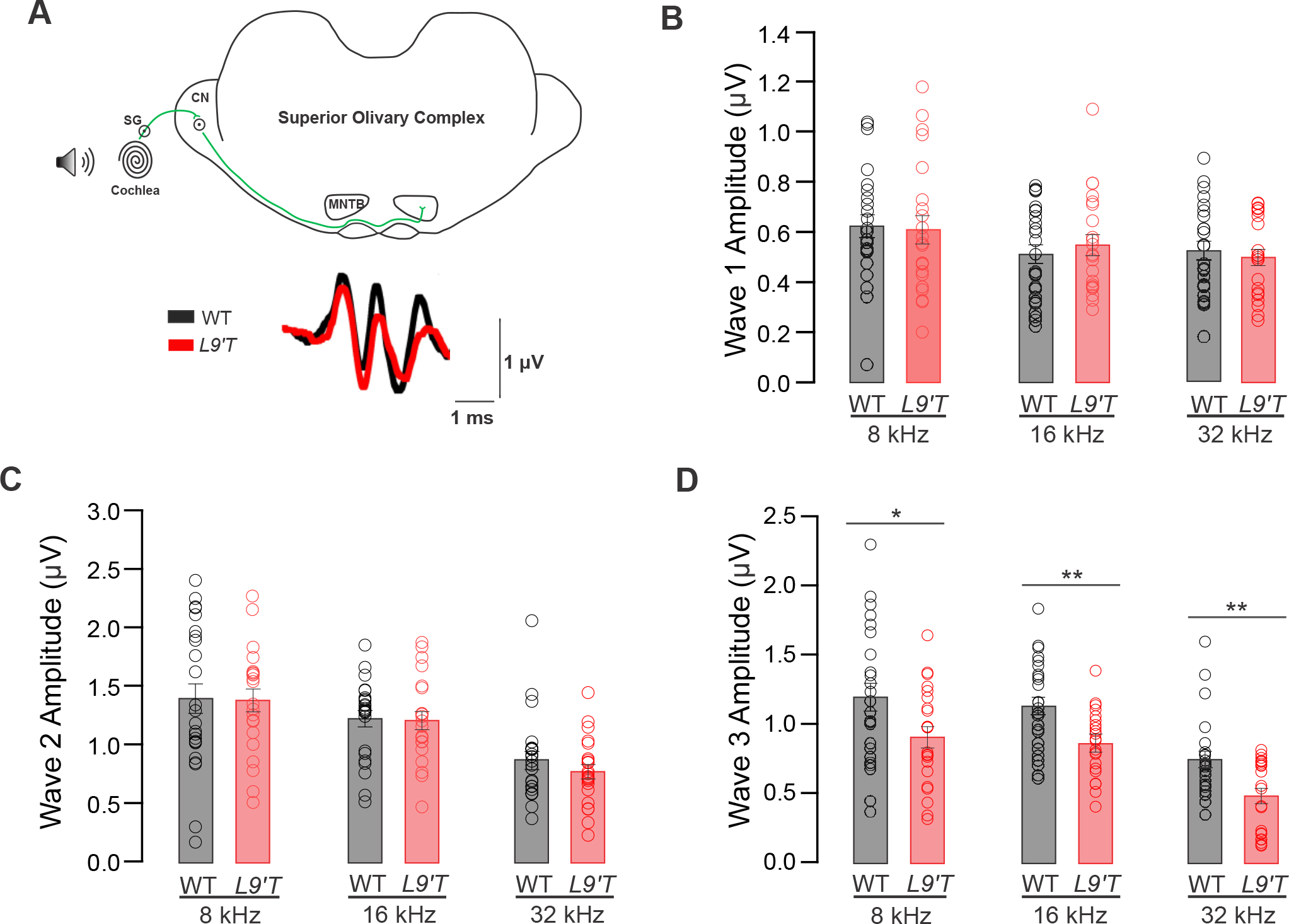
Auditory brainstem response of wave III was smaller in the *L9’T* at P21. **A**. Schematic diagram of the brainstem at the level of the superior olivary complex showing the cochlear stimulation with a speaker. Representative traces of auditory brainstem responses (ABR) of WT (black) and *L9’T* (red) at 80 dB SPL/32 kHz showing waves I, II and III (bottom). ABR wave I (**B**), wave II (**C**) and wave III (**D**) amplitudes at 80 dB SPL for 8, 16 and 32 kHz. No differences were found at all frequencies tested neither for wave I nor wave II. However, *L9’T* mice have lower wave III amplitudes at all the frequencies tested (ANOVA, p<0.05). Bars represent the media ± SEM. All measurements were made at P21. See also Table 1.

### Impairment of synaptic transmission at the calyx of Held in *L9’T* mice

The reduction of ABR peak III amplitude in the *L9’T* mice temporally close to the onset of hearing, raises the question if synaptic transmission at the MNTB level is altered in mice with enhanced MOC activity. To address this point, we performed synaptic studies on slices containing the glutamatergic MNTB-calyx of Held synapse. The rate and amplitude of mEPSCs (Fig. 2A) were recorded in the presence of TTX (1 μM). No differences were observed in mEPSCs amplitude distributions in *L’9T* compared to WT (Fig. 2B.i), even when comparing the mean amplitudes along the medio-lateral axis (WT M: 32.81±2.61 pA, n=8; WT L: 28.37±2.79 pA, n=12; ANOVA, F:1.21, p=0.285; *L’9T* M: 32.82±2.77 pA, n=18; *L’9T* L: 34.28±1.55 pA, n=16; Mann-Whitney test, Z:-0.66; p=0.512, Fig. 2B.ii). However, mEPSC frequency displayed a larger dispersion in *L’9T* mice (Fig. 2C.i), due to a drastically increased mean mEPSC frequency in the lateral region (M: 2.52±0.56 Hz; L: 7.17±1.94 Hz; Mann-Whitney test, Z: -2.11, p=0.035) that was not observed in the WT mice (M: 2.07±0.51 Hz; L: 2.34±0.42 Hz, ANOVA, F:1.16, p=0.689; Fig. 2C.ii). This result suggests that spontaneous transmitter release at the calyx of Held − MNTB synapse is altered in *L9’T* mice.

**Figure 2.**
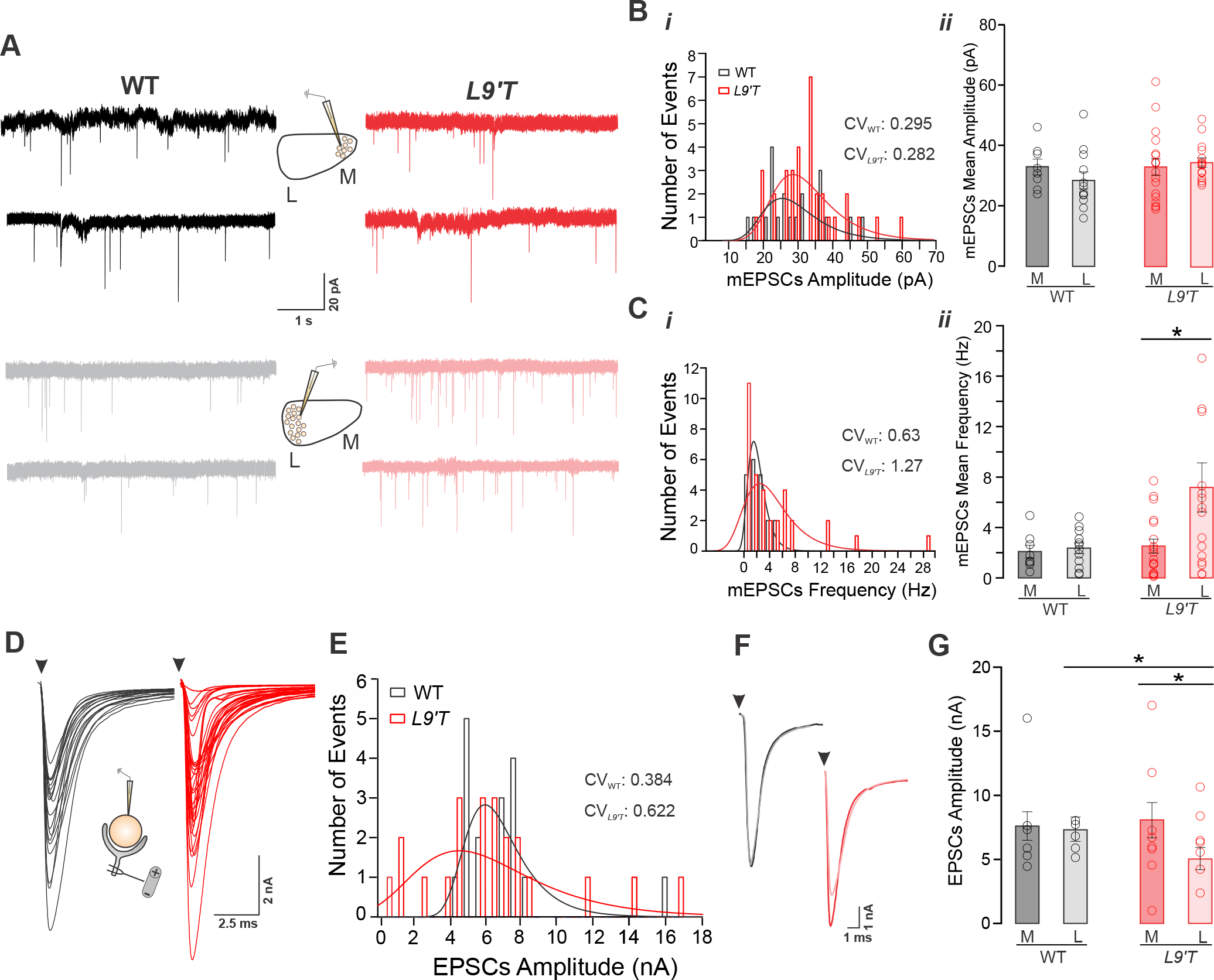
Altered Synaptic Transmission in *L9’T* mice. **A.** Representative traces of miniature excitatory postsynaptic currents (mEPSCs) for WT M (black), WT L (grey), *L9’T* M (red) and *L9’T* L (pink). Examples of two different cells are shown for both genotypes. **B. i.**Histograms for mEPSCs amplitude displaying a similar data distribution for WT and *L9’T*. **B. ii.**The mean mEPSCs amplitude for all conditions displayed no significant differences (WT M: n=8, WT L: n=12, ANOVA, F: 1.21; p=0.285; *L9’T* M: n=18, *L9’T* L: n=16, Mann-Whitney test, Z: −0.66, p=0.512). **C. i**. Data distribution for mEPSC frequency comparing WT and *L9’T* genotypes. The fitting with gaussian function evidences a bias to higher frequencies in the *L9’T*. **C. ii.**mEPSCs frequency was similar for medial and lateral cells in WT (ANOVA, F:1.16, p=0.689). However it was drastically increased in the *L9’T* lateral region (Mann-Whitney test, Z:-2.11; p=0.035). **D.**Representative traces of evoked excitatory post synaptic currents (EPSCs) of MNTB principal cells for WT (black) and *L9’T* (red)**. E.**Histogram distribution of EPSC amplitudes for both genotypes. Note that in this case, histogram fitting with a gaussian curve evidenced a larger tail to lower amplitudes in the *L9’T* mice. **F.**Averaged traces of EPSCs for WT M (black), WT L (grey), *L9’T* M (red) and *L9’T* L (pink). **G.**Mean EPSC amplitude was similar along the tonotopic map in the WT (M: n=9, L: n=10, ANOVA, F: 0.027, p=0.87) but not in the L9’T, where the amplitude in the lateral side decreased (M: n=11, L: n=12, ANOVA, F: 5.07, p=0.0357). Bars represent the media ± SEM.

Principal neurons of the MNTB receive synaptic input from a single giant calyx terminal that generates the stereotyped calyceal EPSC response, which is independent of stimulus intensity above threshold (Fig. 2D). A broader EPSC amplitude distribution in the *L9’T* compared to WT mice was observed (Fig. 2E). Thus, while no significant differences in the unitary medial and lateral EPSC amplitudes were recorded in WT (M: 7.59±1.12 nA, n=9; L: 7.35±0.95 nA, n=10, ANOVA, F:0.027, p=0.87), the evoked synaptic currents in the lateral side (5.07±0.87 nA, n=12) of *L9’T* mice were smaller compared to those of the medial side (8.05±1.37 nA, n=11; ANOVA, F:5.07, p=0.0357, Fig. 2G). Together these results suggest that both spontaneous and evoked synaptic transmission are impaired in the *L9’T* mice, with the lateral low frequency region being the most affected.

### Immature synaptic responses in *L9’T* principal cells

During early development, presynaptic calyx of Held terminals make multiple small contacts on MNTB neurons. This is followed by an early stage of functional and structural transformation (Kandler and Friauf, 1993; Taschenberger et al., 2002; Wimmer et al., 2006; Rodriguez-Contreras et al., 2008), in which multiple inputs strengthen and compete until a final single innervation is established (Holcomb et al., 2013). These multiple early contacts elicit “small”-amplitude glutamatergic currents from each axon (Rodriguez-Contreras et al., 2008; Fig. 3A, left), until a mature calyceal response is established displaying a large, electrical stimulus-independent response (Fig. 3A, right). Thus, the proportion of “small” *versus* “large” responses are reduced with maturation (Rodriguez-Contreras et al., 2008). Given that both “large” and “small” excitatory inputs to MNTB can be distinguished by their electrophysiological profile, the prevalence of small currents in both genotypes was analyzed (Fig. 3B). The probability to find “small” inputs in WT mice was 10% (4 of 35 cells) at P12-P14, whereas this proportion was three-fold larger in *L9’T* mice reaching 31% (12 of 38 cells, Chi-squared test, χ^2^=4.32, df=1, p=0.019).

**Figure 3.**
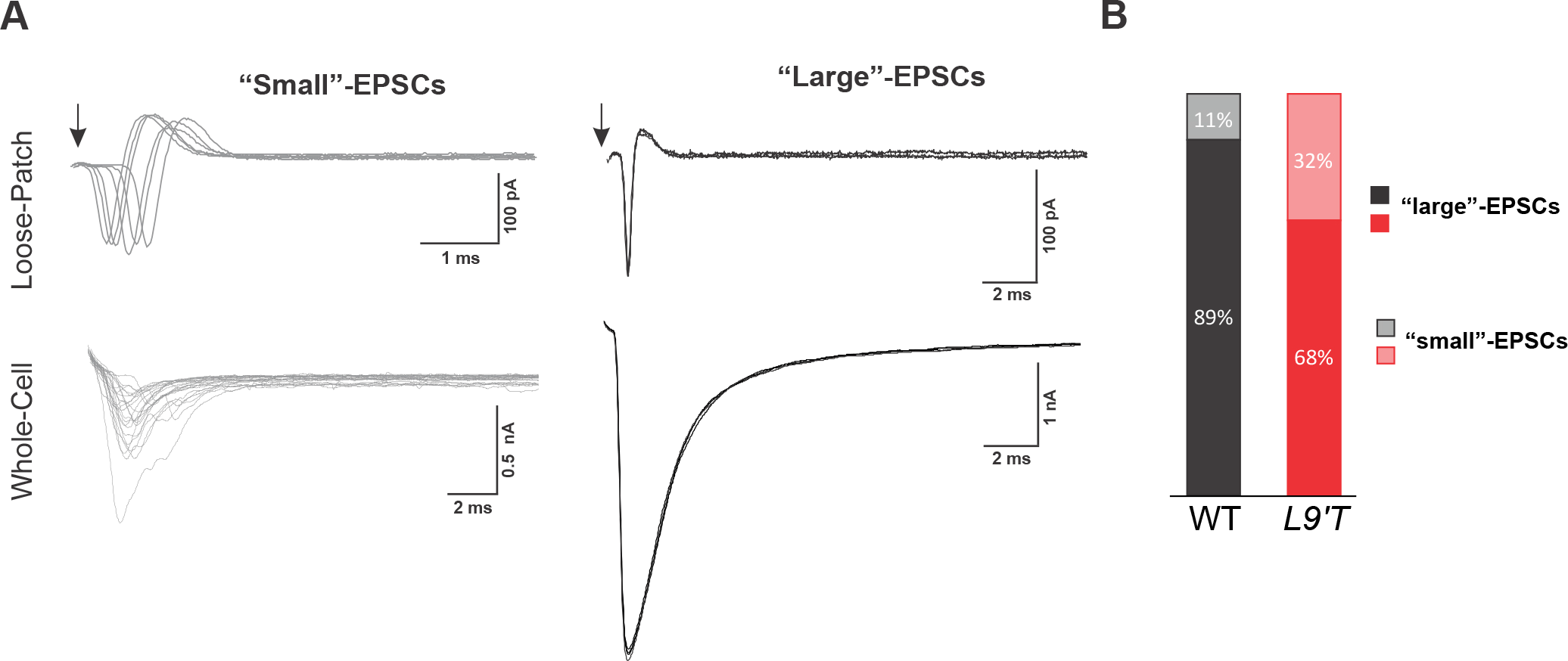
Small-EPSCs were observed more frequently in *L9’T* mice. **A**. Representative traces of two types of EPSCs evoked in MNTB principal neurons by stimulation of the trapezoid body. A large EPSC coming from a calyceal terminal (left) and small-EPCSs from multiple terminals (right). Arrows indicate the position of the stimulation artifact. Cell-attached recordings (loose-patch configuration, upper traces) show action potential currents on MNTB cells. EPSCs were recorded in whole cell configuration (lower traces). Note that calyceal EPSCs did not increase but small-EPSCs amplitude increased with higher stimulus intensity. **B.**A larger proportion of small amplitude connections was observed in *L9’T* mice. In WT mice small-EPSCs were 10% (4 of 35 cells) whereas this proportion was three-fold larger in the *L9’T* mice (31% 12 of 38 cells; Chi-squared test, χ^2^=4.32, df=1, p=0.019).

It has been previously demonstrated that there is a close correlation between the “small” EPSCs and morphological features such as the multiple innervation of MNTB neurons in an off-target manner (Rodriguez-Contreras et al., 2008). In order to analyze these immature features of MNTB innervation, the presence of axon branching was studied by labeling axons with a dextran dye (Fig. 4A). In WT pups (between P12-P14), only canonical unitary calyx terminals contacting a single MNTB neuron were observed (20 slices of 60 μm-thickness, from 5 animals), in accordance with previous descriptions for that developmental period (Rodriguez-Contreras et al., 2008). However, in *L9’T* mice, the presence of axon branching and collaterals were observed (7 of 25 slices from 5 animals; Fig. 4B). Moreover, MNTB principal neurons innervated by at least two presynaptic terminals were observed in *L9’T* mice (2 of 25 slices of 60 μm-thickness from 5 animals; Fig. 4C and D). These last observations are in agreement with electrophysiological results confirming the presence of “multi”-synaptic contacts in the *L9’T* model (see Fig. 3B), in a time window where this feature should be absent or considerably reduced.

**Figure 4.**
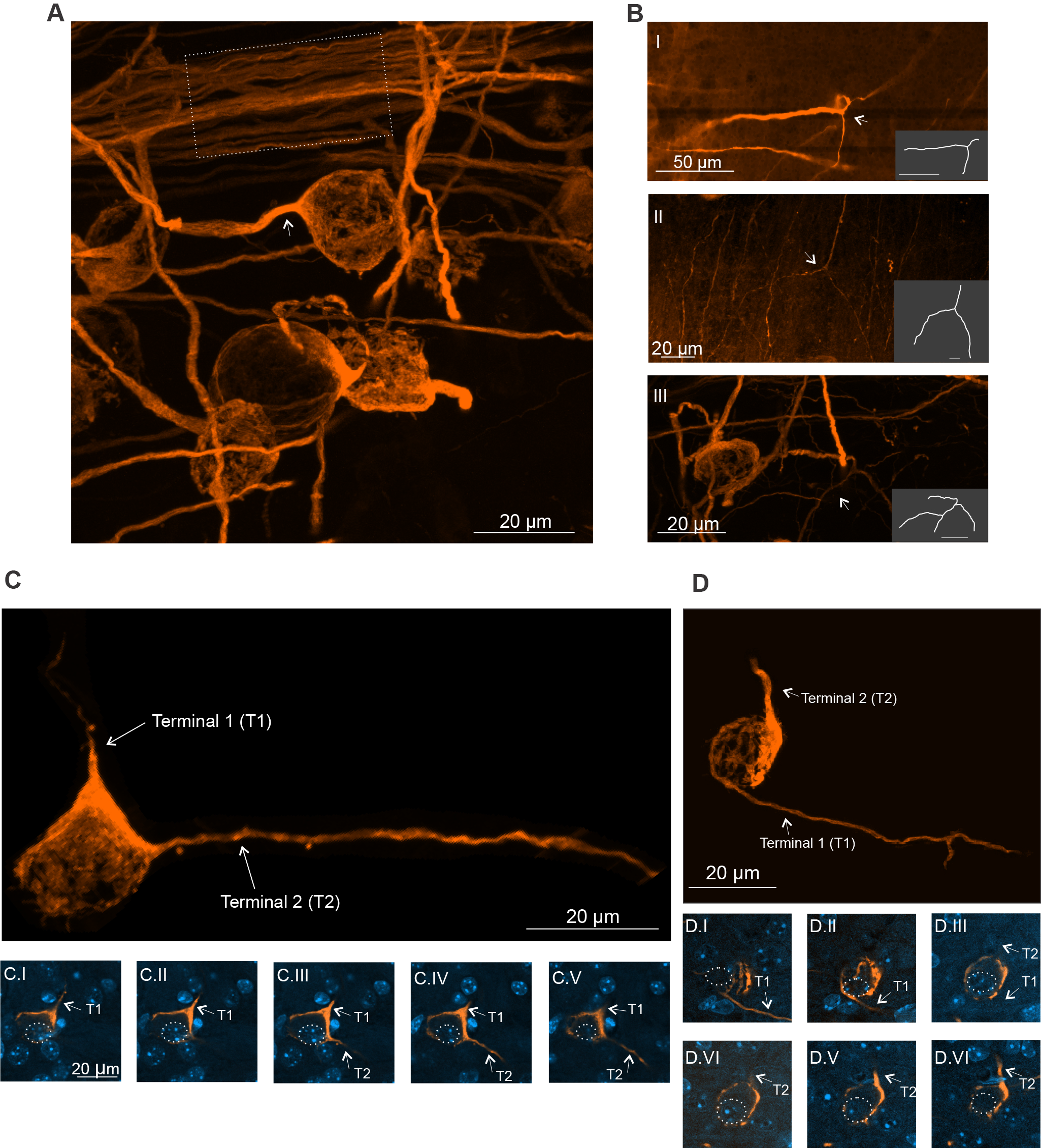
Multiple innervation in *L9’T* MNTB principal cells suggest an impairment during development. Rhodamine-dextran applied *in-vivo* was used to label calyceal terminals. **A.**General overview from the MNTB of an *L9’T* mouse (63X) displaying the normal architecture as seen in WT of the labeled axons (dashed square) and calyx terminals (arrow). **B.**Examples of branching of calyceal afferents visualized in *L9’T* animals. Grey insets display a digitized tracing for each axonal profile. Snapshots were acquired at different magnifications (I: 20X; II: 40X and III: 63X). Arrows indicate the branching point in every case. **C-D.**Examples of two axonal terminals contacting one principal MNTB neuron (indicated by a dash white circle). **C**. Complete Z-stack projection of 17 confocal sections (magnification: 40X). Five representative sections (every 1.14 μm) are shown (**C.I**to **C.V**). Note that insets I to V were slightly rotated to the right respect to image B. **C**. Z-stack projection of 52 confocal sections (magnification: 63X; steps: 0.5 μm). Six selected sections (every 1.5 μm) are displayed (**D.I**to **D.VI**). Arrows indicate axonal terminals. Slices were mounted with medium containing DAPI for nucleus identification (blue label).

### Altered topographic organization of MNTB action potential waveforms in *L9’T* mice

Figures 1 – 4 present evidence for both synaptic and connectivity impairments between calyx of Held terminals and MNTB principal cells in *L9’T* mice. In order to analyse if these alterations were also accompanied by the disruption of the MNTB topographic arrangement, the medio-lateral gradient of different biophysical neuronal properties was analysed (Fig. 5A).

**Figure 5.**
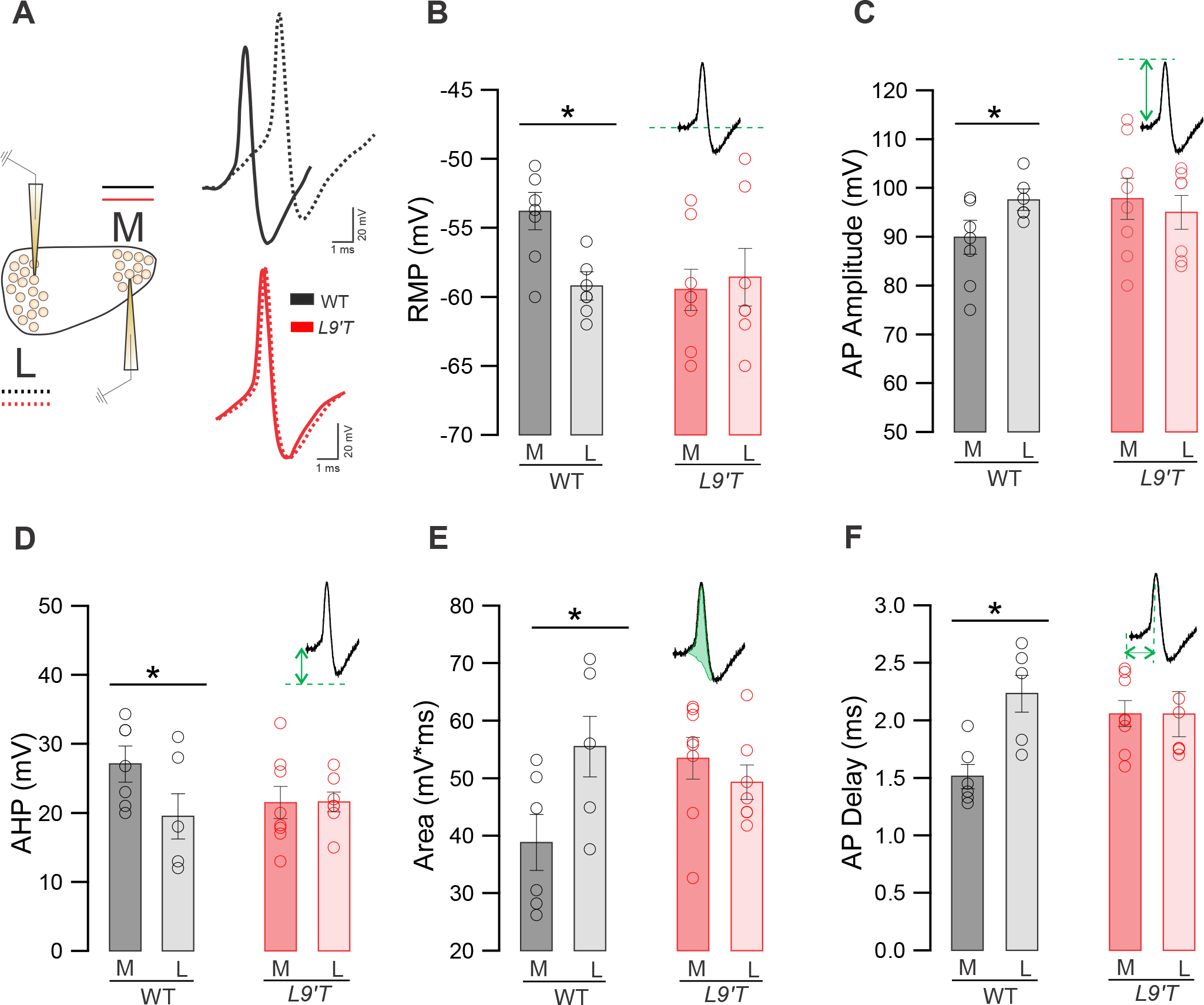
AP shape differed along the tonotopic axis in WT but not in *L9’T* mice. **A**. Superimposed action Potential (AP) waveform of WT (black) and *L9’T* (red) MNTB cells. Filled lines represent the medial and dashed lines the lateral side. **B**. The resting membrane potential (RMP) was more depolarized for medial than lateral cells in WT mice (M: n=7; L: n=6, ANOVA, F: 9.33, p=0.0013). However, no medio-lateral differences were detected in *L9’T* (M: n=8; L: n=7, ANOVA, F: 0.136, p=0.72). AP features were quantified: AP amplitude (**C**), After hyper polarization (AHP; **D**), area (**E**) and AP delay (**F**). Note that in all cases, mediolateral differences in WT AP waveform were abolished in the *L9’T* mice. Bars represent the media ± SEM. See additional information on Table 2.

In the first place we analysed the neuronal resting properties. In WT mice, the resting membrane potential (RMP) was more depolarized for medial (−54.27±0.47 mV, n=7) than lateral cells (−59.49±0.36 mV, n=6; ANOVA, F:9.33, p=0.0013) (Fig. 5B). However, this difference was absent in the *L9’T* mouse model (M: −59.57±0.53 mV, n=8; L: - 58.57±0.79 mV, n=7; ANOVA, F:0.136, p=0.72).

Like other neurons, MNTB principal cells can elicit an action potential (AP) after a positive current injection (Fig. 5A), displaying waveform changes along the tonotopic map (Leao et al., 2006). MNTB cells in WT presented a topographic distribution in AP amplitudes (L>M, 8.6% Fig. 5C), after-hyperpolarization (AHP; M>L, 27.9% Fig. 5D), area (L>M, 42.9% Fig. 5E) and delay (L>M, 47.67% Fig. 5F). However, none of these parameters showed medio-lateral differences in *L9’T* mice (Fig. 5C-F and Table 2). These results demonstrate that, contrary to WT mice, at the MNTB level neither the RMP nor the AP waveform of MNTB principal cells show a medio-lateral differential distribution in mice with enhanced MOC activity.

**Table 2.** AP features. ^1^p = 0.038, ^2^p = 0.045, ^3^p = 0.042; ^4^p = 0.0037. One-way ANOVA

|  | WT |  | *L9'T* |  |
| --- | --- | --- | --- | --- |
|  | M | L | M | L |
| **Amplitude (mV)** | 89.88±1.41 (n=7) | 97.61±0.98 (n=6)<sup>1</sup> | 97.75±1.49 (n=8) | 95.01±1.31 (n=7) |
| **AHP (mV)** | 27.05±1.07 (n=7) | 19.50±1.54 (n=8)<sup>2</sup> | 21.5±0.82 (n=8) | 21.57±0.54 (n=7) |
| **Area (mV\*ms)** | 38.83±1.99 (n=6) | 55.49±2.14 (n=6)<sup>3</sup> | 53.48±1.29 (n=8) | 49.29±1.14 (n=7) |
| **Delay (ms)** | 1.51±0.04 (n=6) | 2.23±0.07 (n=6)<sup>4</sup> | 2.06±0.04 (n=8) | 2.05±0.07 (n=7) |

<sup>1</sup>F: 5.518, p=0.038; <sup>2</sup>F: 4.678, p=0.045; <sup>3</sup>F: 5.418, p=0.042; <sup>4</sup>F: 14.185, p=0.0037

### Lack of medio-lateral gradient of potassium current amplitudes in *L9’T* mice

Hyperpolarization-activated cyclic nucleotide-gated (HCN) channels, which flux both Na^+^ and K^+^, are responsible for the inward I_h_ rectifying current which is present in many auditory neurons and can influence neuronal excitability (Bal and Oertel, 2000). The combination of I_h_ and potassium conductances allow the accurate relay of high frequency auditory information across MNTB neurons and thus the fidelity by which timing information is computed (Hooper et al., 2002; Barnes-Davies et al., 2004; Hassfurth et al., 2009; Mathews et al., 2010; Karcz et al., 2011; Khurana et al., 2012; Baumann et al., 2013). The latero-medial gradient of these conductances was analysed in WT and *L9’T* mice.

Repeated application of negative voltage steps from −50 to −140 mV, displayed an instantaneous (I_I_) followed by a slowly inward (I_s_) current (Fig 6A, inset), leading to a I_h_ (Is-I_I_) current. The specificity of this inward current (mediated by HCN channels) was confirmed by its sensitivity to the selective blocker ZD7288 in all cases (Fig 6A, inset). In agreement with previous observations (Leao et al., 2006), I_h_ currents in WT mice exhibited a medio-lateral gradient (Fig. 6B, −140 mV, M: −372.74±35.48 pA, n=11; L: - 227.21±32.84 pA, n=7, ANOVA F: 7.89, p=0.013). However, in the *L9’T* mice, this difference was absent (M: −381.79±23.95 pA, n=10; L: −332.81±22.27, n=9, ANOVA, F:2.22; p=0.156). Results obtained under voltage-clamp mode were further supported with current-clamp experiments. HCN channels can be activated by a hyperpolarizing current pulse injection which generates a slow “sag” in the membrane potential (Banks et al., 1993; Koch et al., 2004). The sag amplitude in WT medial cells (−51.84±4.13 mV, n=9) was larger compared to that of the lateral region (−39.54±4.78 mV, n=10, ANOVA, F: 5.03; p=0.038). This difference was absent in *L9’T* mice (M: −56.86±6.74 mV, n=7; L: −45±43 mV, n=8; ANOVA, F: 2.05; p=0.17).

**Figure 6.**
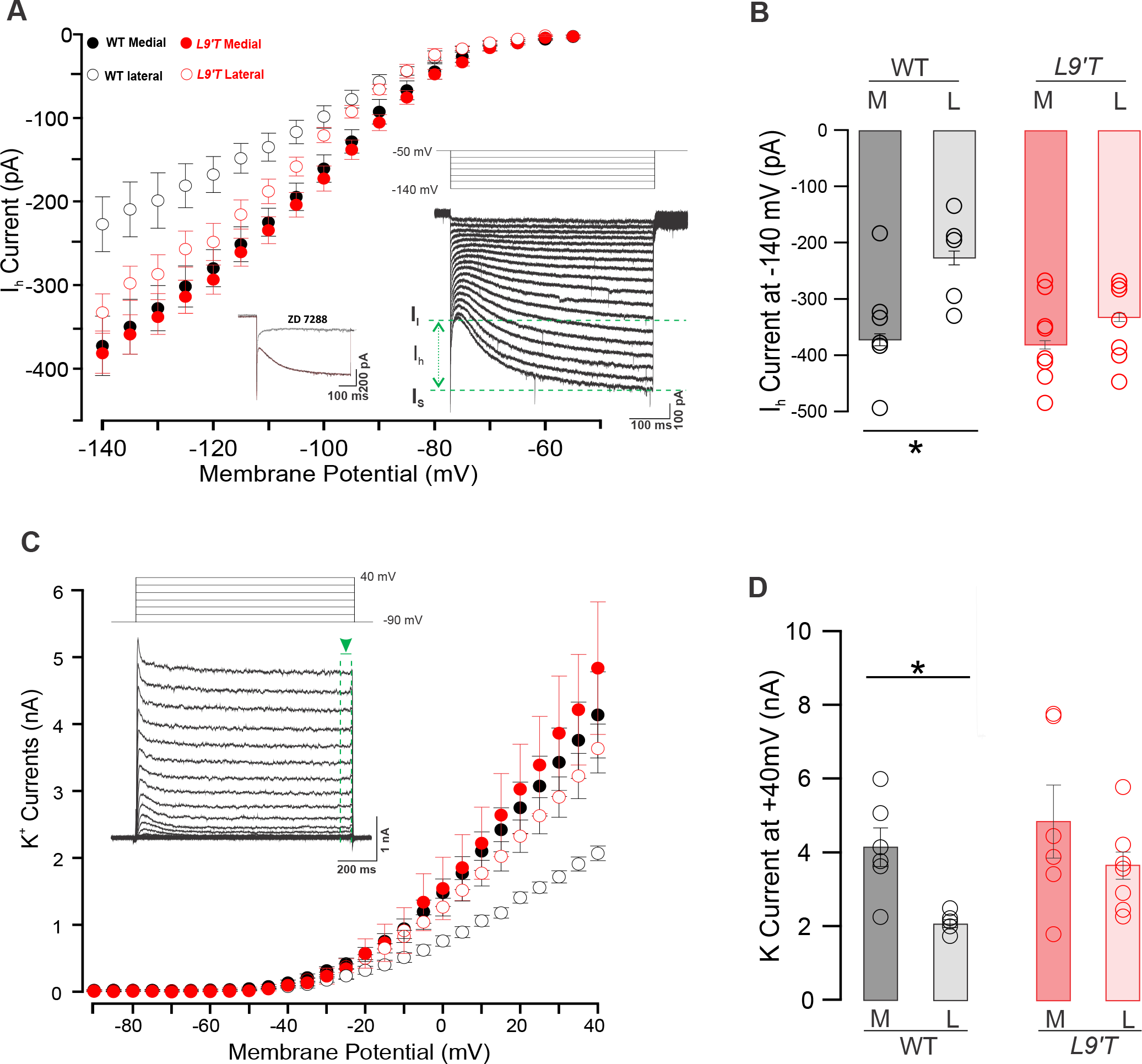
Medio-lateral gradient of potassium currents was absent in *L9’T* mice. **A**. I-V relationship for I_h_ currents exhibited a medio-lateral gradient in WT but this topographic difference disappeared in *L9’T* mice. Bars represent the media ± SEM. **Inset**. Representative traces of I_h_ currents (WT medial) elicited by hyperpolarizing voltage steps from -50 to −140 mV every 10 s for 0.5 s (right). Current responses consisted of an instantaneous inward current (I_I_) and a slowly developing inward current (I_S_). I_h_ amplitudes were determined as I_S_-I_I_. Representative I_h_ trace at −140 mV after bath perfusion with the specific HCN channel blocker ZD 7288 (50 μM, left). **B**. Quantification of maximal I_h_ current amplitude at −140 mV for WT (M: n=11; L: n=7, ANOVA, F: 7.89, p=0.013) and *L9’T* mice (M: n=10; L: n=9, ANOVA, F: 2.22; p=0.156). **C**. I-V relationship displayed a larger potassium current in medial compared to lateral cells in WT mice, but without a tonotopic distribution in the *L9’T* mice (bars represent the media ± SEM). **Inset**. Representative traces at different depolarizing voltage steps (from −90 to +40mV; 5 mV increment during 1s) in presence of TTX (3 μM), CdCl_2_ (50 μM) and ZD7288 (50 μM). Potassium currents were measured at the steady-state (within the last 50 ms, green dashed lines). **D.**Maximal K^+^ current amplitude at +40 mV exhibits a medio-lateral gradient for WT (M: n=6; L: n=6; Mann-Whitney test, Z: 2.72, p=0.0065) but not in the *L9’T* mice (M: n=6; L: n=7, Mann-Whitney test, Z: 1.01, p=0.32).

Potassium conductance was elicited in response to 1 s steps from −90 mV to +40 mV in the presence of CdCl_2_ (50 μM), TTX (3 μM) and ZD7288 (50 μM) to block voltage-dependent calcium, sodium and I_h_ currents, respectively. Outward currents displayed a fast-small inactivating component and a large delayed non-inactivating component (Fig. 6C, inset). Current-voltage relationships displayed topographic differences in WT mice. Thus, at +40 mV the medial MNTB cells exhibited a larger (4.15±0.65 nA, n=6) and the lateral a lower (2.07±0.11 nA, n=6; Mann-Whitney test, Z:2.72, p=0.0065) amplitude of potassium currents (Fig. 6D, inset). In contrast, medio-lateral differences were not observed in *L9’T* mice (M: 4.83±0.99 nA, n=6; L: 3.63±0.37 nA, n=7; Mann-Whitney test, Z:1.01, p=0.32). In summary, potassium currents are not topographically distributed in the MNTB of *L9’T* mice.

## Discussion

Making use of genetically modified mice with a mutation in the α9 nAChR, rendering increased suppression of IHC spontaneous activity (Taranda et al., 2009; Wedemeyer et al., 2018), we show functional and morphological alterations at the glutamatergic MNTB-calyx of Held synapse during the developmental critical period. Lack of latero-medial gradients in several functional properties of the MNTB were observed in mutant mice, together with synapse dysfunction. These alterations remained after hearing onset, reflected in overall reductions of ABR wave III amplitudes.

It has been suggested that the origin of IHC spontaneous spiking activity is located in the Kölliker’s organ that periodically triggers waves of ATP (Tritsch et al., 2007), leading to depolarization and firing of developing IHCs (Tritsch et al., 2010b; Wang et al., 2015). Alternatively, spontaneous activity is intrinsically generated by hair cells, and ATP modulates the firing pattern or coordinates the activity of adjacent cells (Johnson et al., 2011; 2012). A tight regulation of the pattern of IHC action potentials appears as a key feature for auditory development (Jonson et al., 2011; 2013; Sendin et al., 2014). Patterned spiking activity is propagated to spiral ganglion cells (Jones et al., 2007; Trisch et al., 2010a; b), brainstem auditory nuclei (Lippe et al., 1995; Sonntag et al., 2009; Trisch et al., 2010a; Clause et al., 2014) and auditory cortex (Babola et al., 2018). In cochlear explants, both the application of extracellular ACh, as well as the electrical stimulation of efferent terminals, inhibits action potential generation through the activation of α9α10 nAChRs coupled to SK2 channels present in IHCs (Glowatzki and Fuchs, 2000; Goutman et al., 2005; Gomez-Casati et al., 2009). Johnson et al. (2011) have suggested that ACh released from efferent terminals is essential for setting a bursting firing pattern in apical IHCs. Moreover, Sendin et al. (2014) have shown that pharmacological block of α9α10 nicotinic receptors elicits an increase in the IHC spontaneous discharge rate. Similarly, a higher firing rate of developing MNTB neurons has been observed in α9 knock-out mice (Clause et al., 2014). Since MNTB spontaneous activity is originated in the cochlea (Trisch et al., 2010b), the latter observations most likely arise from the lack of MOC innervation in these mutants. However, the role of α9α10-mediated cholinergic transmission to IHCs in modulating spontaneous spiking activity has been challenged (Tritsch et al., 2010a). The results shown in the present study clearly indicate that the transient α9α10-mediated transmission to IHCs is crucial for modulating spiking-dependent development of the auditory system. Since during development a modest stimulation rate of efferent fibers is sufficient to produce strong, near-maximal inhibition of IHC firing (Moglie et al., 2018), and this inhibition is exacerbated in the *L9’T* mutant mice (Wedemeyer et al., 2018), the resultant phenotype most likely results from the silencing of IHC spiking activity.

### Synaptic transmission

In the absence of calcium channels responsible for the release of glutamate by IHC (Ca_V_1.3 knock-out mice), development of synaptic transmission at the calyx of Held is impaired (Erazo-Fischer et al., 2007). Although these mice lack auditory nerve activity, those experiments have not clearly distinguished between spontaneous and sound-evoked afferent spiking. The reduction observed in wave III amplitude in the present work can derive from alterations in synaptic transmission at the calyx of Held in *L9’T* mice, since this peak of the ABR is dependent upon synchronous synaptic activity at this nucleus. The increased number of asynchronic “small”-EPSCs compared to the calyceal-monosynaptic synchronic stimulus intensity-independent EPSCs in mutant mice, can account for this phenotypic observation. These asynchronic “small”-EPSCs probably result from the lack of synaptic refinement observed in *L9’T* mutant mice, a dynamic process that normally takes place around P2-P4, when calyceal collaterals are pruned, gradually disappear and principal cells end up being contacted by a single calyx of Held (Rodríguez-Contreras et al., 2008; Hoffpauir et al., 2006, 2010; Holcomb et al., 2013). It is worth noting that the high proportion of multi presynaptic contacts of principal MNTB neurons observed in *L9’T* mutants differs from the apparent normal morphology observed in α9 knock-out mice (Clause et al., 2014). The mechanisms leading to the persistence of immature calyces in mutant mice is unknown, but most likely reside on the lack of cues needed to strengthen competing synaptic inputs to a mature innervation as described for the neuromuscular junction (Wu et al., 2010), the climbing fiber innervation of Purkinje cells (Watanabe and Kano, 2011), and the retinal ganglion cell innervation of the dorsal lateral geniculate nucleus (Hong and Chen, 2011). In this regard, the *L9’T* mutant phenotype resembles that of the bone morphogenetic protein conditional knock-out, with impaired nerve terminal growth, loss of mono-innervation and less mature transmitter release properties (Xiao et al., 2013). It has been reported that developmental pruning of calyceal collaterals is independent of sound-evoked activity (Rodríguez-Contreras et al., 2008). The present results suggest that it is highly dependent upon the transient MOC efferent innervations that tightly controls spontaneous spiking activity.

The increased frequency of mEPSCs in mutant mice might derive from the higher number of cells contacted by multiple calyces. Thus, individual active zones from different axons could independently contribute to spontaneous release, leading to an enhancement of mEPSC frequency. Alternatively, changes in the expression of proteins involved in vesicle docking, priming, as well as Ca^2+^ sensors and other SNARE-binding proteins (Schneggenburger and Rosenmund, 2015), might lead to the same observation. Since evoked synaptic currents were only recorded from principal neurons contacted by a single calyx, the reduction in the amplitude of EPSCs indicates that even in “mature” synapses, synaptic transmission is altered in *L9’T* mutants. Moreover, the present results show that the intrinsic passive and active properties of MNTB cells which are established during the early developmental period (Hoffpauir et al., 2006), are also altered in *L9’T* mutant mice, suggesting upstream alterations in synaptic transmission from the MNTB to lateral SOC, an inhibitory pathway in the mammalian sound localization system (Kandler, 2004).

### Tonotopy

Characteristic frequencies of neuronal response to acoustic stimulation are tonotopically arranged. This is the result of a precise topography of connections that is preserved along the auditory pathway (Friauf and Lohmann,1999; Rubel and Fritzsch, 2002). This tonotopy is maintained and/or reflected in the topographic arrangement of neuronal functional properties (Li et al., 2001; Barnes-Davies et al. 2004; von Hehn et al. 2004; Brew & Forsythe, 2005; Pienkowski and Harrison, 2005; Leao et al., 2006). One key question is how tonotopy is shaped by either spontaneous sound-independent or sound-evoked auditory nerve activity (Rubel & Fritzsch, 2002; Sanes and Bao, 2009). Animal models have been used in order to dissect the effect of spontaneous activity in the establishment of auditory topography. For example, congenitally *dn/dn* deaf mice exhibit alterations in the structural and functional topographic arrangement of several auditory nuclei, indicating disrupted tonotopy (Leao et al., 2006). Although initially considered as a model of disrupted spontaneous spiking activity (Durham et al.,1989), recent findings challenge this hypothesis. Thus, in *dn/dn* which bear a deletion in the transmembrane channel-like protein (TMC) 1 (Kurima et al., 2002), prehearing IHCs develop normally and fire spontaneous calcium action potentials (Marcotti et al., 2006).

Experiments performed in α9 knock-out mice, which lack functional transient efferent innervation to IHCs (Vetter et al., 1999; 2007), have shown normal levels of spontaneous activity in the MNTB but altered temporal spiking patterns (Clause et al., 2014). Moreover, although the overall tonotopy is maintained, the strengthening and silencing of inhibitory MNTB-LSO connections before hearing onset is impaired. This resulted in a reduced sharpening of functional topography, as the consequence of a reduction in axonal pruning. In the present work we demonstrate that the enhancement of MOC activity, and therefore the increased inhibition of IHC spontaneous activity (Wedemeyer et al., 2018), disrupts the topographic specificity of several MNTB characteristics. These include the resting membrane potential, action potential waveform and I_h_ and potassium currents. Since these conductances allow accurate auditory transmission at high rates across MNTB neurons (Hooper et al., 2002; Barnes-Davies et al., 2004; Hassfurth et al., 2009; Mathews et al., 2010; Karcz et al., 2011; Khurana et al., 2012; Baumann et al., 2013), one could speculate that *L9’T* mice display frequency discrimination impairments, similar to those reported for α9 knock-out mice (Clause et al., 2017). One could propose that the transient MOC efferent system is a first checkpoint of auditory spontaneous activity. It is interesting to note that the dysfunction of this first checkpoint of IHC activity provided by MOC efferents (as exhibited in α9 knock-out and *L9’T* knock-in mice) is not compensated by homeostatic mechanisms, such as described for *Vglut3* knock-out mice which lack IHC glutamatergic synaptic transmission (Babola et al., 2018). This might indicate that spontaneous activity in the auditory system is always dictated by the cochlea when IHCs are functionally connected to spiral ganglion neurons. Since central auditory disfunction is observed both in α9 knock-out (Clause et al., 2014) and knock-in mice (present results), the transient MOC efferent transmission to IHC most likely provides the patterning of spiking activity rather than acting as an on/off switch to spontaneous activity.

## Acknowledgements

We thank Lucas G. Vattino and Marcelo J. Moglie for critical reading of the manuscript, Juan D. Goutman for fruitful discussion and assistance with data analysis through routines implemented in Igor Pro. We also thank Claudia Gatto for excellent technical assistance. This work was supported by Agencia Nacional de Promoción Científica y Tecnológica, Argentina (A.B.E.), and NIH Grant R01 DC001508 (Paul A. Fuchs and A.B.E.). MNDG is the recipient of The Company of Biologist travelling fellowship (DEVTF-160703) and a “Bec.Ar” fellowship for a short stay supported by the Argentinian government.

